# Influence of organs, body size and growth on domoic acid depuration in the king scallop, *Pecten maximus*

**DOI:** 10.64898/2026.03.23.708139

**Authors:** Eline Le Moan, Hélène Hégaret, Margot Deléglise, Marie Ambroziak, Jean Vanmarldergem, Amélie Derrien, Aourégan Terre-Terrillon, Florian Breton, Caroline Fabioux, Fred Jean, Jonathan Flye-Sainte-Marie

## Abstract

Since 1995, European fisheries of *Pecten maximus* faced the presence of *Pseudo-nitzschia* species, which are able to produce the neurotoxin domoic acid responsible for Amnesic Shellfish Poisoning (ASP). As filter-feeders, scallops can accumulate and retain domoic acid much longer than most of the other bivalves, from months to years. When concentrations exceed the regulatory threshold, fisheries are closed leading to economic crisis. Inter-individual variability increases the difficulty to predict the depuration dynamics. Quantifying the correlations between domoic acid depuration in *P. maximus* and individual physiological traits, particularly body size, could improve the understanding of contamination and depuration. In this study, toxin dynamics in organs were analysed and the effects of body size and growth were assessed. This analysis was based on two datasets, one experimental and one in situ, of depuration monitoring of *P. maximus* exposed to a natural bloom of toxic *P. australis*. Results show that the distribution of domoic acid shifted among organs between the contamination and after two months of depuration. Toxin concentrations negatively correlate with body size during contamination and after two months of depuration, but shift to a positive correlation after 7 months of depuration. This shift suggests that the smaller scallops accumulate more domoic acid and depurate it faster. Thus, dilution by growth can explain the reversal of the correlation between domoic acid and body size throughout depuration. These results yield useful information for modelling such mechanisms, providing valuable tools for scallop fishery management facing ASP.

**Graphical Abstract:** 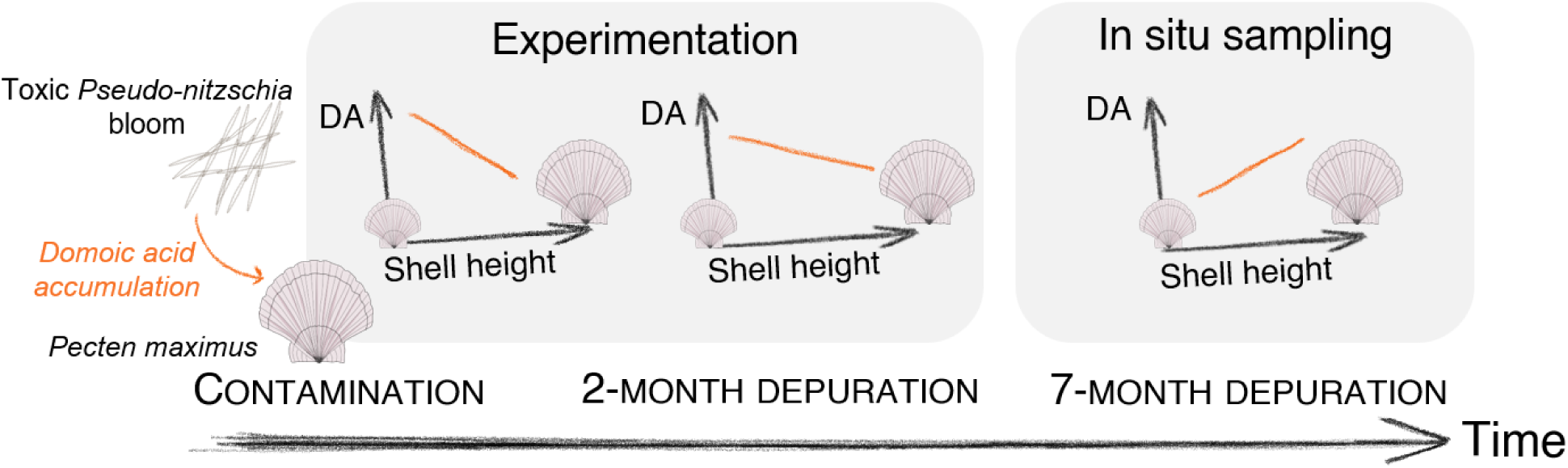

**Highlights:**

- Experimental and in situ datasets allowed to quantify DA proportion dynamics in organs of *P. maximus*
- DA concentration and body size are negatively correlated during contamination phase, but positively correlated after a 7-month depuration
- Considering dilution by growth is important for young scallops to assess DA depuration dy-namics
- Both depuration rate and dilution by growth need to be considered to model DA depuration over the whole scallop size range

## 1 Introduction

Bivalve molluscs, as filter-feeders, are vulnerable to harmful algal blooms (HABs), which are proliferations of microalgal species that may affect the environment notably by producing toxins (Hallegraeff et al., 2004). By accumulating these toxins through feeding, bivalves may convey them through the food web and transfer them to higher trophic levels, like marine mammals and humans (Bejarano et al., 2008). In commercially exploited species, toxin levels are routinely monitored to avoid human intoxication. When the toxin concentrations in bivalve tissues exceed regulatory limits, the fishing or aquaculture activities are temporarily closed (Basti et al., 2018). Recurrent closures, particularly for high-value species have significant socio-economic consequences (FAO, 2025a,b).

Domoic acid is a neurotoxin mainly produced by more than 26 species of the genus *Pseudo-nitzschia* (Bates et al., 2018), an ubiquitous pelagic diatom. It is responsible for the Amnesic Shellfish Poisoning (ASP) symptom, first detected in Canada in 1987, after ingestion of contaminated mussels (Wright et al., 1989) which led to three human deaths and 153 illnesses (Bates et al., 1988). In humans, gastrointestinal symptoms occur within 24 hours, while neurological symptoms, such as memory loss, confusion and coma, emerge after 48 hours (Perl et al., 1990; Krasner et al., 2025). The domoic acid retention by primary consumers depends on both accumulation process through feeding and depuration. The kinetics of these processes depend on the species. Some commercially exploited species, such as the blue mussel *Mytilus edulis* (Mafra et al., 2010), the Chilean scallop *Argopecten purpuratus* (Álvarez et al., 2020), or the giant scallop *Placopecten magellanicus* (Douglas et al., 1997), depurate the toxin relatively quickly, leading to moderate economic consequences. By contrast, other species such as the Pacific razor clam *Siliqua patula* (Horner et al., 1993), and the king scallop *Pecten maximus* (Blanco et al., 2002), retain the toxin for very long periods, up to months or years. The latter, *P. maximus* is able to accumulate high levels of domoic acid compared to other bivalves, as reported by García-Corona (2023). Yet, this high and long retention remains poorly understood. The involved physiological mechanisms are likely based on a lack of membrane transporters that allow rapid excretion of domoic acid (Mauriz and Blanco, 2010) and its trapping in autophagosome-like vesicles (García-Corona et al., 2022) for long periods.

This long toxin retention is one of the challenges faced by the scallop fishery, as it can lead to harvesting bans for several months or years. For instance, Blanco et al. (2002) showed that individuals that were naturally exposed to *Pseudo-nitzschia* took more than one year to fall below the regulatory threshold of 20 *mg kg^−^*^1^. In Galicia (Spain), domoic acid contamination is so persistent that fishing is almost permanently banned, except for selective evisceration (INTECMAR, https://www.intecmar.gal/). In other locations such as Ireland, domoic acid is often detected in scallops. Therefore, the sale of shucked scallops (adductor muscle with or without gonad) is allowed if the domoic acid concentrations remain below 250 *mg kg^−^*^1^ in the whole animal and 20 *mg kg^−^*^1^ in the edible part (Food Safety Authority of Ireland, 2024). In France, domoic acid was first detected in 2004 with contaminated scallops from November 2004 to June 2005 (Amzil et al., 2009), including the whole fishing season from October to March. Since then, contaminations have been detected in some French regions every two years (REPHYTOX, 2023).

To assist fishing managers and fishermen, it is crucial to develop methods to reduce the contamination and/or to accelerate the depuration. Such methods, however, may be unfeasible in real-world situations (Leal and Cristiano, 2026). For example, antioxidants that were tested to accelerate domoic acid depuration in *P. maximus* proved to be unsuitable for fisheries (Vanmaldergem et al., 2023). Alternatively, another approach is to predict the depuration period to anticipate the fishing season, which includes the estimation of the depuration rates (Free and Fang, 2026). To achieve this, understanding the mechanisms behind the prolonged retention of domoic acid in *P. maximus* is crucial. For this purpose studying the source of variability in domoic acid concentration may provide useful insights.

The rapid depuration of domoic acid in *Crassostrea virginica*, *M. edulis* (Mafra et al., 2010) and *A. purpuratus* (Álvarez et al., 2020) was attributed, in part, to toxin transfer from the digestive glands to other organs. In *P. maximus*, Campbell et al. (2001) showed a high inter-individual variability in domoic acid concentrations for a pool of 10 individuals from the west coast of Scotland, with a shell length ranging between 9 and 12 cm. So far, very little is known about the toxicity of individual organs in *P. maximus* and their dynamics. Only Blanco et al. (2002) suggested that the domoic acid transfer between organs was low. Therefore, the quantification of domoic acid in each organ at the peak of contamination and during depuration may offer insights into the depuration mechanisms of *P. maximus*.

The correlation between the individual size and the toxin dynamics has been studied in several bivalve species. In naturally contaminated mussels, Novaczek et al. (1992) showed that smaller mussels accumulated higher domoic acid concentrations in their organs, and depurated faster. Similar patterns were observed in oysters exposed to *P. multiseries*: smaller individuals contained more toxin and exhibited faster depuration rate compared to larger ones (Mafra et al., 2010). The relationships with body size, however, were not constantly observed in mussels during the same experiment. In *P. maximus*, such complex relationship between domoic acid concentration and size has been observed (Bogan et al., 2007). Correlations with body size help in understanding the mechanisms implied in the toxin accumulation and depuration, which are particularly important to model the toxin kinetics. For example, Bogan et al. (2007) highlighted a correlation between food ingestion (including toxins) and weight. To clarify how body size influences domoic acid concentration, the present study examines the total domoic acid concentrations across a broader size range of *P. maximus* than in Bogan et al. (2007).

In addition to the size of the organism, growth may have an important effect on the toxin dilution within the organism, especially considering long retention time. The dilution by growth was estimated at 54% of the total toxin loss for a 2-month period exposition of the surfclam *Spisula solidissima* to paralytic shellfish toxin (Bricelj and Shumway, 1998). The importance of such phenomenon has also been demonstrated for the depuration of ciguatoxin (Bennett and Robertson, 2021). Blanco et al. (2002) proposed that domoic acid may be lost along with biomass, and that biomass reduction does not necessarily lead to an increase of toxin concentration, suggesting that dilution by growth may not be a major factor in *P. maximus*. However, the authors did not account for growth, and the individuals were in suboptimal conditions, as indicated by the observed biomass loss. To improve the estimation of domoic acid depuration duration in *P. maximus*, it is necessary to confirm whether growth-driven dilution is a major factor to incorporate.

The main objective of this work is to assess how the dynamics of domoic acid depuration depend on the individual physiology, addressing the three following questions: (i) Is the proportion of domoic acid similar in each organ over time during the depuration phase? (ii) Is there a relationship between domoic acid concentration and size, and does the pattern change through the depuration period? (iii) Should the dilution by growth be considered in addition to the depuration process and to what extent? To address these questions, this study relies on two different datasets for depuration monitoring, using both experimental and in situ conditions.

## 2 Materials and methods

### 2.1 Scallop samplings

#### A two-month experimental depuration (Dataset 1)

A total of 90 king scallops were dredged at Camaret-sur-mer, France (48*^◦^*31N, 4*^◦^*60W) on 8 April 2021, following a natural bloom of *P. australis* in Brest Bay that occurred in April 2021 REPHY (2023). The animals were transferred to the experimental facility of the Tinduff hatchery (Bay of Brest, France), scrubbed to remove epibionts, and measured (weight and shell height). A total of 30 individuals were sampled and dissected for initial domoic acid measurement and biometry. The remaining organisms were placed on well-aerated sand in 800 *L*-fiberglass tanks, continuously supplied 0.2 *µm*-filtered seawater from the Bay of Brest (16*^◦^C* and salinity of 35), for two months. They were fed with a mix of *Tisochrysis lutea*, *Chaetoceros* sp. and *Skeletonema* sp. at a rate of 10 *×* 10^9^ *cell ind^−^*^1^ *d^−^*^1^. During the experiment, 22 animals died after 2 months, the 38 remaining scallops were sampled and dissected for biometry and domoic acid quantification.

#### A seven-month in situ depuration (Dataset 2)

On 15 January 2025, 244 king scallops were dredged at two locations in Brest Bay during depuration, following a natural bloom of *P. australis* that occurred in April 2024. Both locations were selected based on the zones defined by the French Monitoring programme for Phycotoxins in marine organisms (REPHYTOX, 2023): “039 - Rade de Brest - 039-S-281 Rade de Brest - Nord” where 123 individuals were collected, and “039 - Rade de Brest - 039-S-282 Rade de Brest - Sud” where 121 individuals were dredged. The individuals were transferred to experimental facilities at the laboratory of environmental marine science (LEMAR) and maintained overnight before biometric measurements and organ dissection. The shells were dried in an oven at 60*^◦^C* for 2 days and stored for further shell growth reading, while the total flesh was stored at -20*^◦^C* before analyses.

### 2.2 Biometry measurements

Dataset 1 - The individuals were weighted (total flesh) and measured (shell height) before organ dissection. The digestive gland (containing the stomach and part of the intestine) and the gonad were then separated from the other organs. Each organ or group of organs was weighted and stored at -20*^◦^C* before domoic acid analyses.

**Dataset 2 -** The individuals were weighted (total, shell, flesh) and measured (shell length, height, width).

**Both datasets -** The condition index was calculated as the ratio of flesh wet weight to cubic shell height expressed in *g cm^−^*^3^ (Zeng and Yang, 2021).

### 2.3 Estimation of the dilution by growth factor

Dataset 2 - To make the annual growth striae more visible, the shells were placed in a 10% acetic acid bath for 45 seconds and then rinsed with fresh water to clean them. For each individual, the annual striae were determined and counted under a binocular magnifier. The number of striae was used to determine the age class of the individual. The shell height (from the umbo to the edge of the shell) was measured for each striae as the individual growth between two age classes. Specifically, the growth period of the individual during the toxin depuration phase was considered as the growth season between the two winters (2024 and 2025). It could then be quantified as the difference in shell height between the last observed striae and the total shell height of the individual. To account for the dilution by growth, the increase of individual body-volume between these two winters was estimated. Assuming isomorphy, the volume of an individual is proportional to the cube of its shell height (Kooijman, 2010). Therefore, the ratio of the cubic shell height in the last winter (January 2024) to the cubic shell height at the sampling day 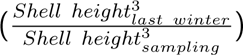 approximates the dilution factor for the toxin. The maximum value of this factor is 1 (no growth), meaning that there is no dilution by growth. If the factor is below 1 (indicating growth), then dilution by growth may occur. The domoic acid concentration is multiplied by this factor to back-calculate the initial concentration before the dilution by growth.

### 2.4 Domoic acid quantification in tissues

#### 2.4.1 Tissue preparation for extraction

Dataset 1 - For each of the 68 individuals in dataset 1, the digestive gland, gonad, and rest of organs were separately ground using Ultraturax before collecting 0.200 *±* 0.005 *g* of each sample. The samples were then stored in a 2-*mL* Eppendorf ©Safe-Lock tube and kept at -20*^◦^C* before extraction and domoic acid quantification.

Dataset 2 - The whole flesh of each of the 243 scallops was ground using Ultraturax, until the grind was completely homogeneous. Then 0.200 *±* 0.005 *g* of each sample was collected in a 2-*mL* Eppendorf ©Safe-Lock tube and stored at -20*^◦^C* prior to extraction.

#### 2.4.2 Tissue extraction

For all 200 *±* 5 *mg* samples, 450 *µL* of a mix Methanol/Water (*MeOH*/*H*_2_*O*) (50/50, v/v) were added, along with 250 to 300 *mg* of 100-250 *µm* glass beads. Then, the samples were ground using the Retsch Mixer Mill MM 400, for 3 *min* at 30 *Hz* then centrifuged at 15000 *g* for 5 *min*, after which the supernatant was isolated and stored. On the pellet, an additional 450 *µL* of the same mix *MeOH*/*H*_2_*O* (50/50, v/v) was added and the steps of grounding and centrifugation were repeated to retrieve the supernatant. The supernatants were combined and the volume was adjusted to 1 *mL* with the mix *MeOH*/*H*_2_*O* (50:50, v:v). Finally, 200 *µL* of this extract were filtered using 0.2 *mm* nylon centrifugal filtering tubes (VWR International, Radnor, PA, USA) at 10000 *g* for 5 *min* at 4*^◦^C*, and the filtrate was stored in amber glass vial 0.3 *mL* at -20*^◦^C* until quantification.

#### 2.4.3 Domoic acid quantification

Domoic acid quantification was performed using liquid chromatography coupled to tandem mass spectrometry (LC-MS/MS), following the method described by Ayache et al. (2020). The analysis was conducted on a Shimadzu UFLCxr system coupled with an API 4000 QTrap hybrid quadrupole mass spectrometer (Sciex, Concord, ON, Canada) equipped with a heated electrospray ionisation (ESI) source. Chromatographic separation was carried out on a reversed-phase Phenomenex Luna Omega C18 column (150 × 2.1 *mm*, 3 *µm*, Phenomenex, Torrance, CA, USA). The separation used a mobile phase composed of aqueous eluent A (100% *H*_2_*O*+0.1% formic acid) and organic eluent B (95% acetonitrile/5% *H*_2_*O* +0.1% formic acid). The gradient transitioned from eluent A to eluent B as follows: 5% at min 0, 18.6% at min 17, 95% at min 17.5, held at 95% until min 19.5, reduced to 5% at min 20, and maintained at 5% until min 25. The flow rate was 200 *µL min^−^*^1^ and the injection volume was 5 *µL*. The column temperature was set at 30*^◦^C*. Domoic acid detection was achieved by multiple reaction monitoring (MRM) in positive ion mode. The five transitions and MS/MS parameters are listed in Supp. Table S1. The 312.1 *≥* 266.1 m/z transition was used for quantification, while the remaining transitions were used for confirmation. Quantification was performed using the Certified Reference Material Domoic Acid (National Research Council of Canada, NRCC) with a 6-point calibration curve. The limit of detection (LD, S/N = 3) and limit of quantification (LQ, S/N = 10) were 0.08 and 0.25 *ng mL^−^*^1^, respectively, corresponding to 0.4 *ng g^−^*^1^ and 1.25 *ng g^−^*^1^ of domoic acid in organ.

For dataset 1, as the domoic acid concentrations were measured per organ, this concentration was multiplied by the wet weight of each organ to get the quantity per organ. The total domoic acid quantity in the organism is the sum of these quantities, which was then divided by the total flesh wet weight to obtain the total domoic acid concentration in the whole organism.

#### 2.4.4 Domoic acid depuration rate estimation

The domoic acid depuration rate was estimated by fitting an exponential decay model using a non-linear least squares regression defined as: [*DA*]*_t_* = [*DA*]_0_ *× exp*(*−τ ×* (*t − t*_0_)), with [*DA*]_0_ the domoic acid concentration at initial time, *τ* the depuration rate, *t* the time at which the domoic acid concentration was calculated and *t*_0_ the initial time. For dataset 1, the depuration rates were estimated for the total flesh and for each organ separately. For dataset 2, depuration rates were estimated for the total flesh in both zones. The observations from this study were combined with the observations from the REPHYTOX monitoring programme, between June 2024 and December 2024 (REPHYTOX, 2023). This national programme quantifies phycotoxin in exploited shellfish species using a pool of 10 individuals above the minimal commercial length. The “north zone” from this study was combined with the “039 - Rade de Brest - 039-S-281 Rade de Brest - Nord” REPHYTOX location, and the “south zone” was combined with the “039 - Rade de Brest - 039-S-282 Rade de Brest - Sud” zone of the REPHYTOX monitoring programme.

#### 2.4.5 Back-calculation of domoic acid concentrations

For dataset 2, based on the domoic acid concentrations in January 2025 and the depuration rate estimated for this dataset, the domoic acid concentration observed in June 2024 was back-calculated. In order to assess whether depuration rate only, dilution by growth only, or both processes are important to consider in the domoic acid concentration decrease in scallops, three scenarios were tested: (1) dilution by growth only, by applying the dilution by growth coefficient of each individual and considering a depuration rate of 0 *d^−^*^1^, (2) depuration only, by applying the exponential decay model transformed to calculate the initial concentration 7 months before with the mean estimated depuration rate per zone, and (3) both processes, by applying the dilution by growth coefficient of each individual and the mean estimated depuration rate per zone. The depuration rate used for back-calculation was the one estimated previously for each location “Rade de Brest - Nord” and “Rade de Brest - Sud” (as detailed in section 2.4.4). The outcome of each of the three scenarios was applied to the individual domoic acid concentrations measured in January 2025 to back-calculate the potential concentration in June 2024. The general pattern of the resulting concentrations depending on shell height was first assessed. Then, for individuals above the commercial size, the average of the back-calculated concentrations was compared to the observed one reported in the REPHYTOX programme: 148.8 *mg kg^−^*^1^ in the North and 129.6 *mg kg^−^*^1^ in the South.

#### 2.4.6 Statistical analyses

Both datasets - All analyses were performed using R software v4.4.1 (R core team, 2022), and the results are presented as mean ± standard deviation. Normality was not achieved on the residuals (Shapiro test), thus Wilcoxon tests were performed to assess the significance of the differences between times of the experiment (dataset 1) and between zones (dataset 2). Correlations between variables were assessed using Spearman correlation coefficients using the “cor” function of the “stat” package, and were represented as a correlation matrix using the “ggcorrplot” function in the “ggcorrplot” package. The conditions of application of each test was performed on residuals thanks to the “performance” package.

Dataset 1 - The relationships between domoic acid concentration, shell height and their interaction with time (initial and final time of the experiment) were assessed per organ and for total flesh using linear models (“lm” function). For both the gonad and muscle+mantle, homoscedasticity was not validated on the residuals, therefore, a linear model with permutation was used (the “lmp” function in the “lmPerm” package). To test the contribution of shell height, time, and their interaction, a type III ANOVA was performed (the “Anova” function in the “car” package).

Dataset 2 - The method used to estimate the depuration rate based on observations is described in Le Moan et al. (2025). Briefly, an exponential decay model is fitted to the data (see section 2.4.4), thanks to a non-linear least squares regression using the “nls” function. The initial value was the first measurement of the domoic acid concentration in June 2024 in the REPHYTOX monitoring programme (REPHYTOX, 2023). Linear models were computed on the domoic acid concentrations according to shell height for (i) measured data in January 2025 and log transformed data to meet the condition of linearity of the model, and (ii) back-calculated toxin concentrations using the three scenarios.

## 3 Results

### 3.1 A two-month experimental depuration (Dataset 1)

#### 3.1.1 Temporal dynamics of physiological traits and domoic acid concentration Biometry

The wet weight of total flesh (**Figure 1A**, Wilcoxon test: p = 0.5) and of muscle+mantle (**Figure 1D**, p = 0.81), as well as the shell height (**Figure 1E**, p = 0.37) were not significantly different between the initial and final times. The final weight of the digestive gland (**Figure 1B**, p = 0.003) and of the gonad (**Figure 1C**, p = 10*^−^*^9^), and the condition index (**Figure 1F**, p = 10*^−^*^14^), however, were significantly lower than those measured at the initial time. At the initial time, the digestive gland, gonad and muscle+mantle represented 11 ± 2%, 14 ± 2% and 76 ± 3% of the total flesh weight, respectively. At the final time, these proportions significantly changed (Wilcoxon tests, p < 0.05) to 9 *±* 1%, 6 *±* 2% and 85 *±* 3%, respectively.

**Figure 1:**
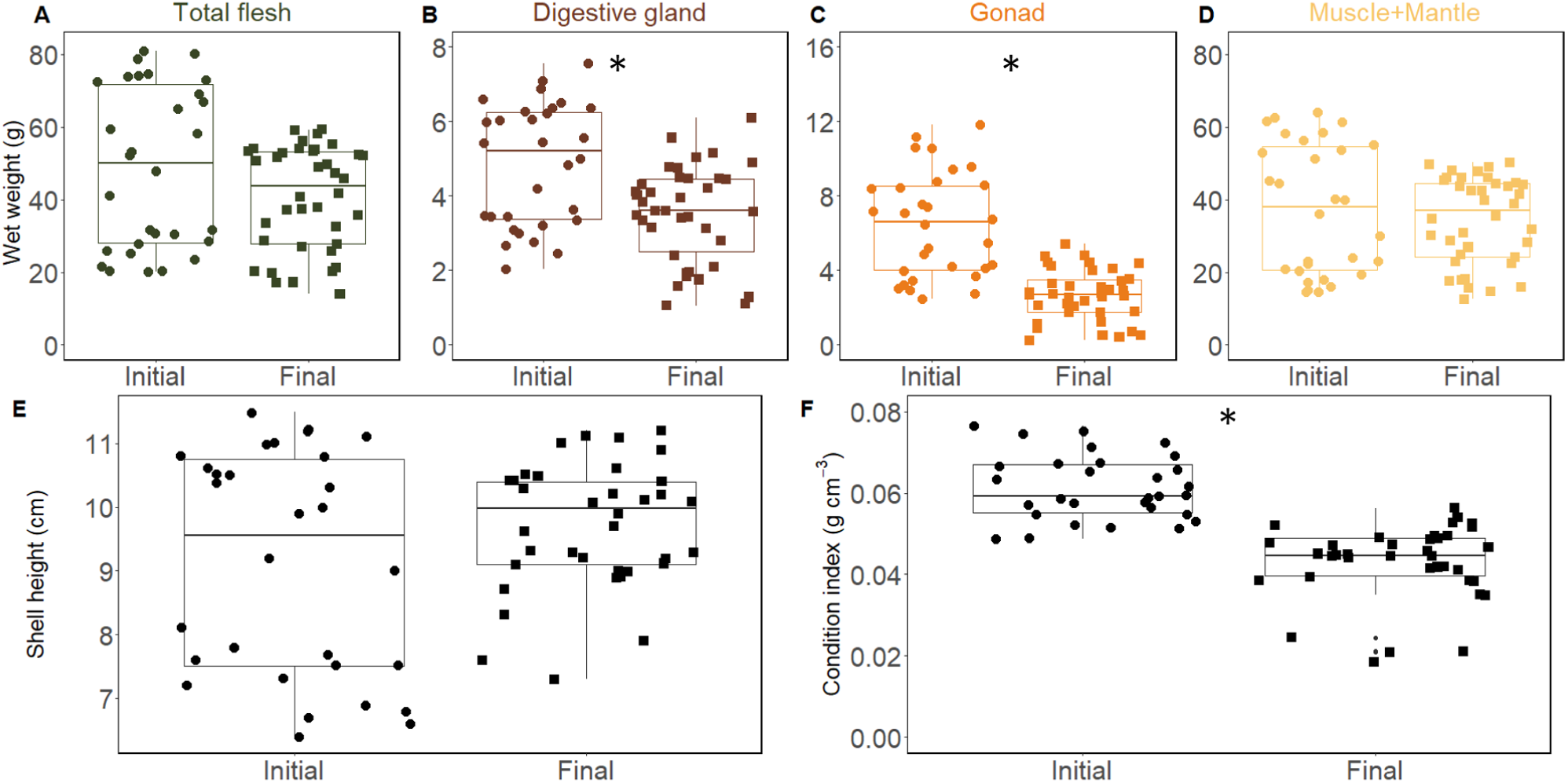
Biometry of the individuals, A) total flesh wet weight (*g*), B) digestive gland wet weight (*g*), C) gonad wet weight (*g*), muscle and mantle wet weight (*g*), E) shell height (*cm*) and F) condition index (flesh wet weight/shell height^3^, in *g cm^−^*^3^) at initial and final time. Significance (p < 0.05) derived from the Wilcoxon test is given with an asterisk.

Domoic acid concentrations

All individuals were contaminated with domoic acid at concentrations higher than the regulatory threshold of 20 *mg DA kg^−^*^1^ (**Figure 2 A-D**). The domoic acid concentrations (mean *±* SD) were significantly greater (Wilcoxon tests, p < 0.001) at initial time compared to final time for digestive gland (initial: 1829 *±* 304, final: 994 *±* 267 *mg kg^−^*^1^), muscle+mantle (initial: 4.47 *±* 2.47, final: 2.1 *±* 0.97 *mg kg^−^*^1^) and total flesh (initial: 200.8 *±* 57.3, final: 97.3 *±* 56.1 *mg kg^−^*^1^), but lower for gonads (initial; 5 *±* 2.9, final: 8.3 *±* 5.7 *mg kg^−^*^1^). The estimated depuration rates were 0.014 *d^−^*^1^ for total flesh, 0.010 *d^−^*^1^ for the digestive gland, 0.013 *d^−^*^1^ for the muscle+mantle. The increase in toxin concentration in the gonads was estimated at 0.008 *d^−^*^1^.

**Figure 2:**
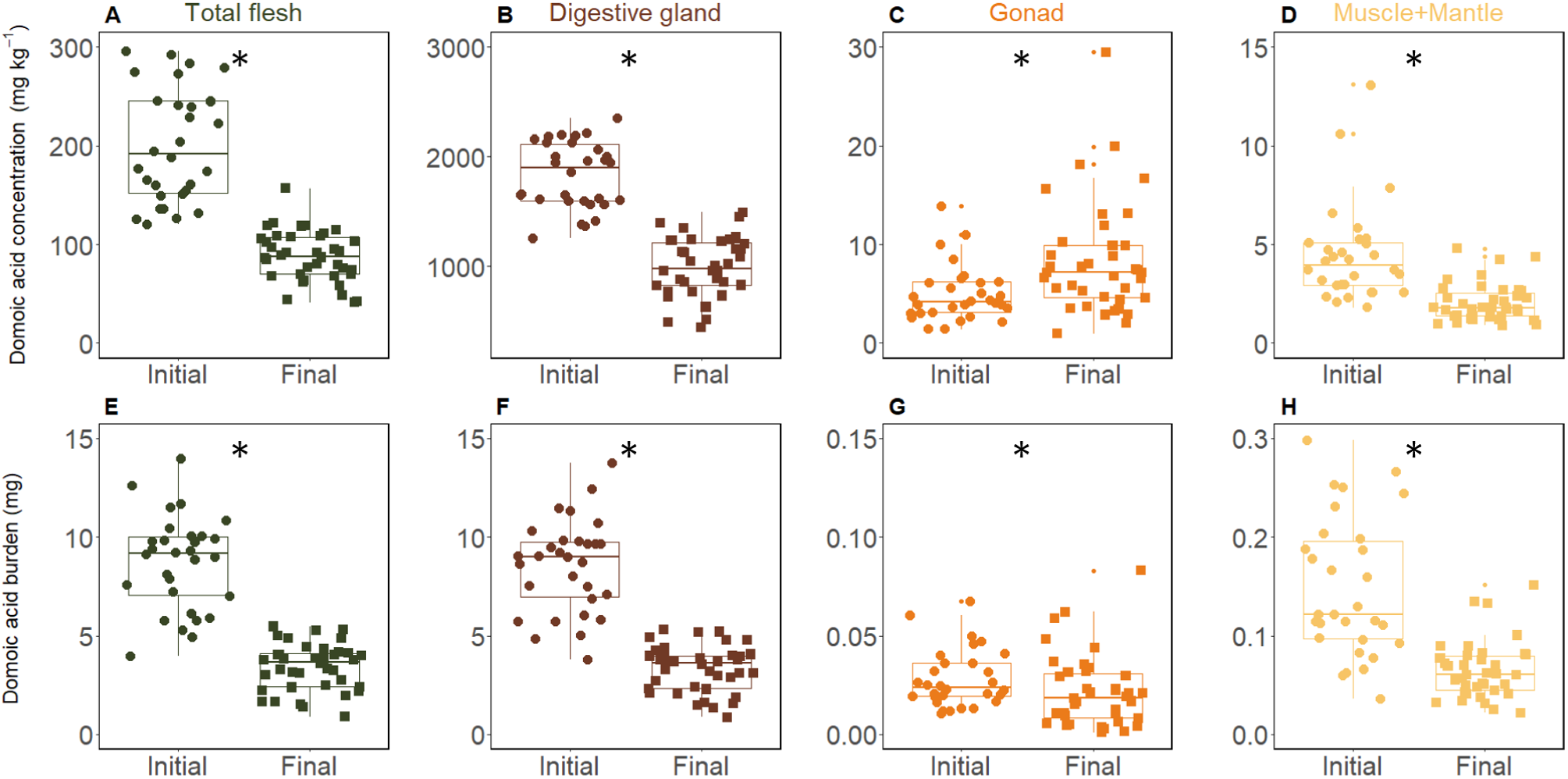
Domoic acid (DA) concentration (*mg kg^−^*^1^) (A-D) and burden (*mg*) (E-H) in total flesh (A,E) and per organ: digestive gland (B, F), gonad (C, G), muscle+mantle (D,H), for initial and final time of the experiment. The little dots represent the outliers from the boxplot. Significance (p < 0.05) from the Wilcoxon test is given with an asterisk.

Domoic acid burden

The domoic acid burden (**Figure 2 E-H**) was significantly greater (Wilcoxon tests, p < 0.001) at the initial time compared to the final time for total flesh (**Figure 2E**, initial: 8.7 *±* 2.4, final: 3.4 *±* 1.2 *mg*) and for all organs. A two-fold decrease was observed for the digestive gland (**Figure 2F**, initial: 8.7 *±* 2.4, final: 3.4 *±* 1.2 *mg*) and the muscle+mantle (**Figure 2H**, initial: 0.15 *±* 0.07, final: 0.07 *±* 0.03 *mg*). The decrease of domoic acid burden in the gonads was minor (**Figure 2G**, initial: 0.03 *±* 0.01, final: 0.02 *±* 0.02 *mg*, p = 0.04).

#### 3.1.2 Proportion of domoic acid quantity in tissues

The proportion of domoic acid differed between the organs (**Figure 3**). The digestive gland contained 97% of the total domoic acid amount, with a significant but small decrease over time (Wilcoxon test, p = 0.008). Then, the muscle+mantle contained 2% of the total domoic acid quantity that did not vary through time (p = 0.14), and the proportion in the gonads increased from 0.4% to 0.6% (p = 0.008) at the final time.

**Figure 3:**
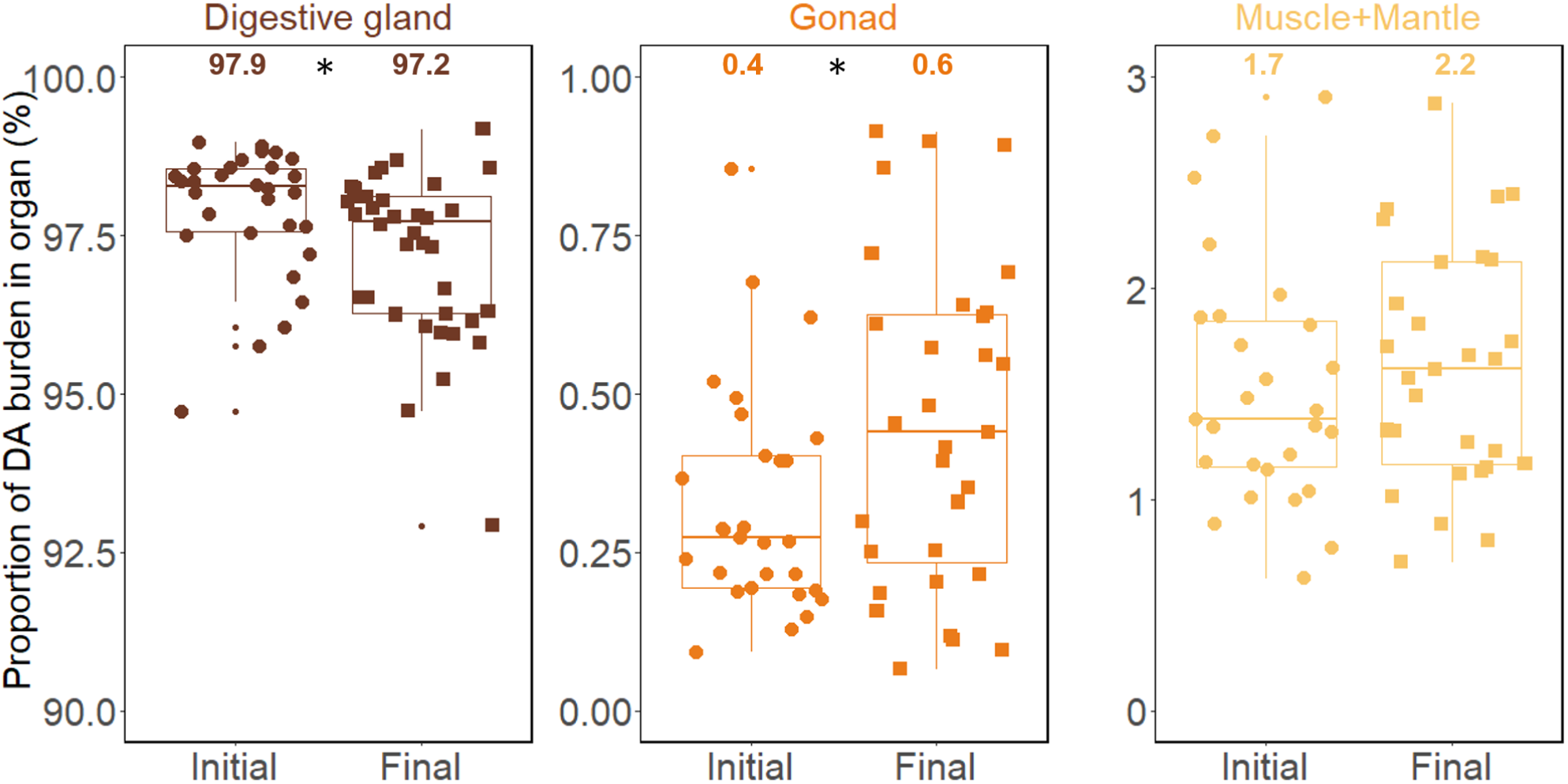
Proportion (%) of domoic acid (DA) burden in the digestive gland, gonad, and muscle+mantle, relative to total domoic acid content at initial and final time of the experiment. The mean is given above the boxplot and significance (p < 0.05) from the Wilcoxon test is given with an asterisk.

#### 3.1.3 Domoic acid concentrations per organ and correlations with body size

At initial time (**Figure 4A**), the shell height and weights (flesh and per organ) were negatively correlated with domoic acid concentrations (Spearman coefficients below -0.44 and -0.42 respectively). Domoic acid concentrations in total flesh and in digestive gland, however, were positively correlated with condition index (0.57 and 0.46 respectively). The correlations were not significant for domoic acid concentrations in muscle+mantle and in gonads (blank squares, **Figure 4A**).

**Figure 4:**
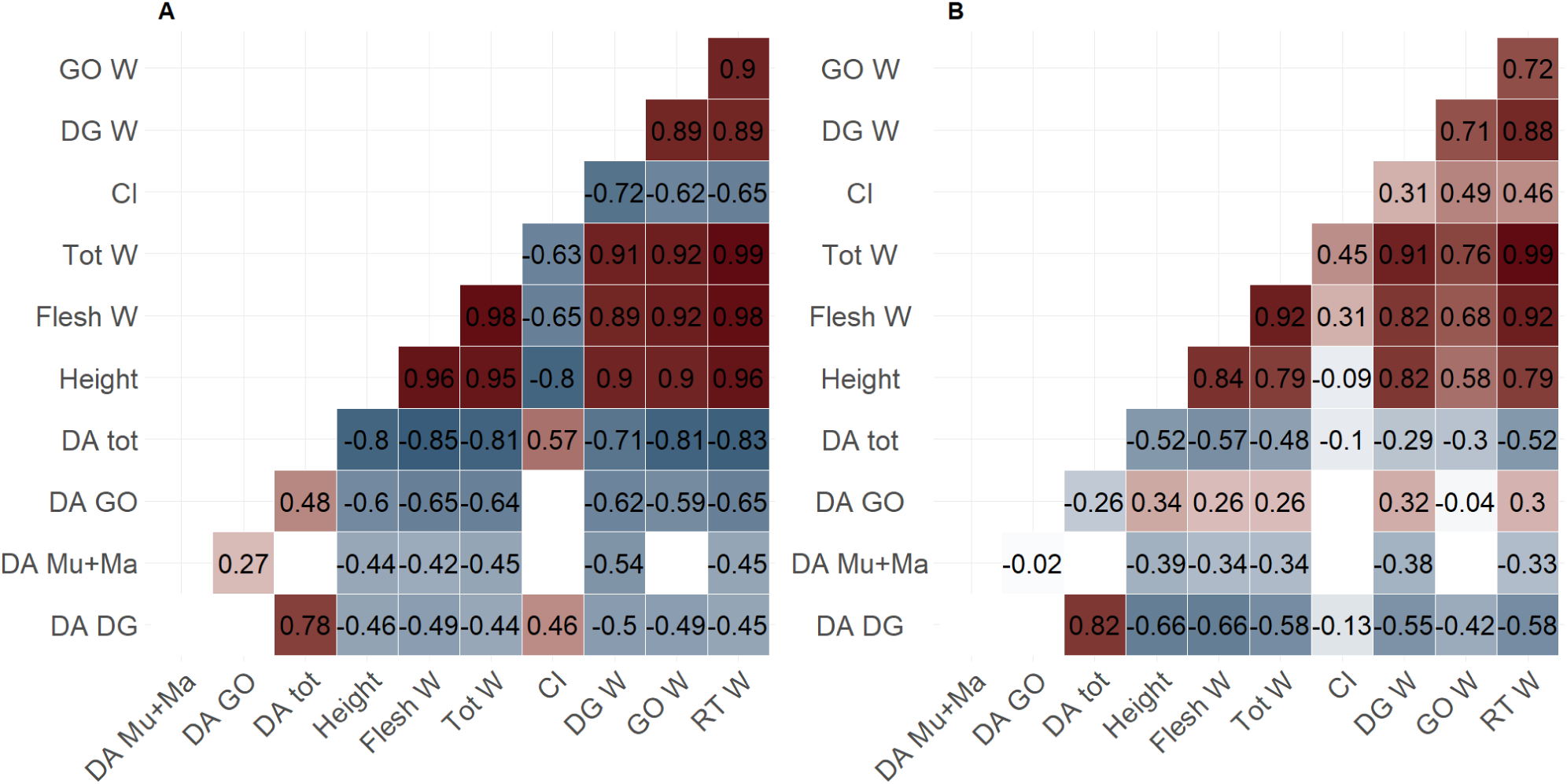
Matrix of Spearman correlation coefficients for (A) initial and (B) final time. The coefficients are given on the matrix (colour and number), no significant correlations are left blank. The labels correspond to the tissues: GO (gonad), DG (digestive gland), Mu+Ma (muscle+mantle), RT (rest of organs), Tot (total flesh); and the variables: W (wet weight), CI (condition index), DA (domoic acid concentration).

At final time (**Figure 4B**), the shell height and weights (total and per organ) were negatively correlated with domoic acid concentrations in muscle+mantle (-0.39 and -0.34 respectively), in digestive gland (-0.66 for both measurements), and in total flesh (-0.52 and -0.57 respectively), but they were positively correlated with gonad (0.34 and 0.26 respectively). The domoic acid concentrations in total flesh and digestive gland were still significantly correlated with the condition index (coefficient of -0.1). At both times, domoic acid concentration in the digestive gland was highly correlated with total domoic acid concentration (initial: 0.78, final: 0.82). Because shell height and flesh wet weight are highly correlated (> 0.8), shell height was used to compare domoic acid concentrations in the subsequent results.

Strong correlations existed between shell height and domoic acid concentrations in total flesh but also per organ (**Figure 5**). Globally, when the shell height is large, the domoic acid concentration is low (**Figure 5A,B,D**), except for gonads at the final time (**Figure 5C**). The domoic acid concentration in the total flesh (**Figure 5A**) was significantly lower when the shell height was larger (type III ANOVA, p = 0.0018). In addition, there were effects of the time (p = 10*^−^*^5^) and of the interaction between shell height and time (p = 0.014). Consequently, the relationship between concentration and shell height was stronger at initial time than at final time. The slopes between initial and final times significantly differed. The domoic acid concentration in the digestive gland (**Figure 5B**) was negatively correlated with shell height (p = 10*^−^*^5^), and this relationship was similar through time with no time effect and no interaction between time and shell height (p > 0.05). There was no general relationship between domoic acid concentration in the gonad and shell height (**Figure 5C**, p = 0.16); the concentrations were significantly correlated with the shell height at initial time, but not at final time, with a positive slope. However, there was a significant effect of time (p = 0.006) and between time and shell height (p = 0.0009). The domoic acid concentration in the mantle+muscle **(Figure 5D**) was significantly lower when shell height was larger (p = 0.0027). There was an effect of the time (p = 10*^−^*^6^), but no significant interaction between time and shell height (p = 0.6).

**Figure 5:**
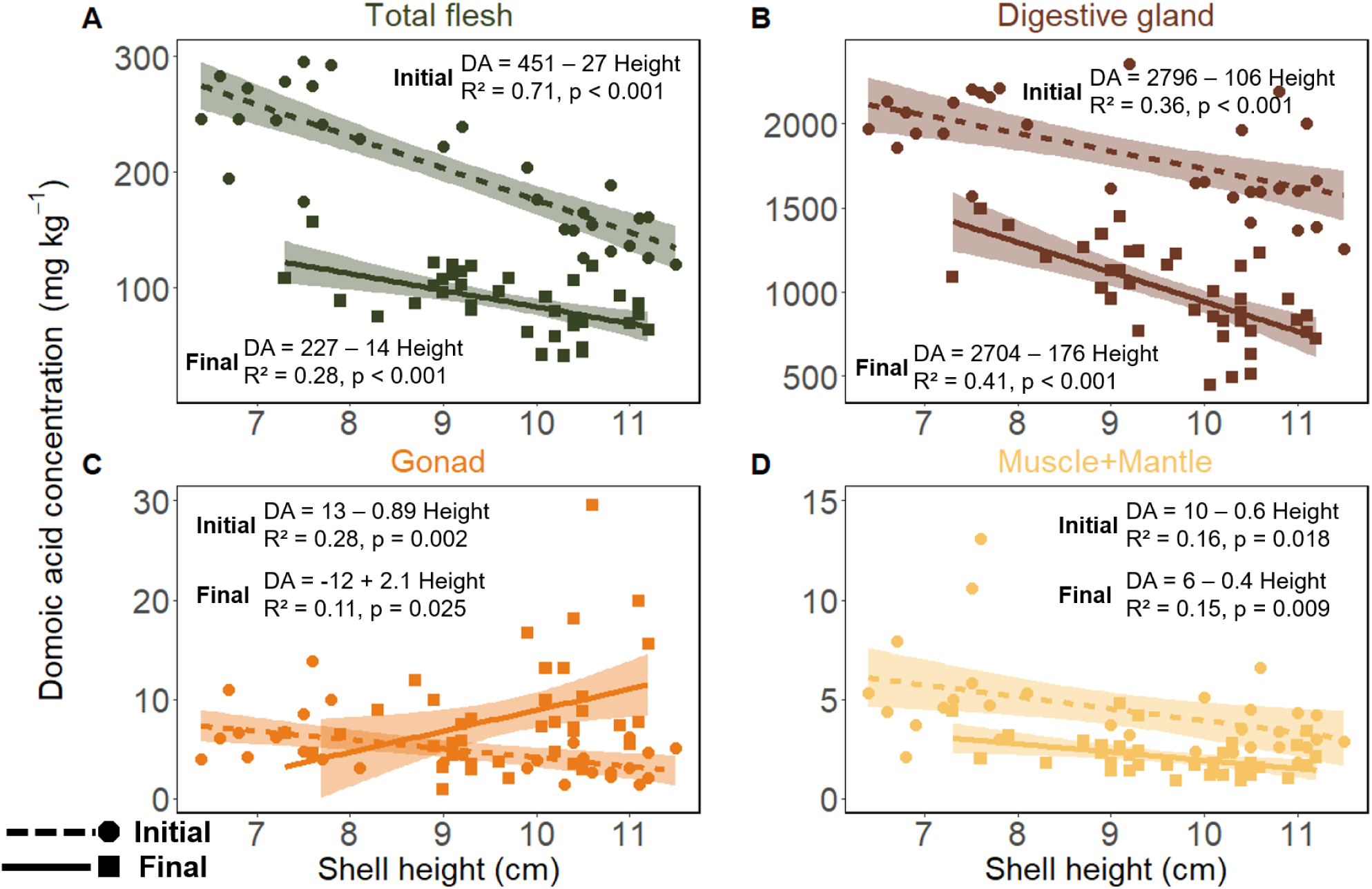
Domoic acid (DA) concentration (*mg kg^−^*^1^) according to shell height (*cm*), for 30 individuals at initial time (circles and dashed line) sampled during the contamination by *P. australis* and for 38 individuals at final time (squares and plain line) sampled after 60 days of depuration. Concentration in A) total flesh, B) digestive gland, C) gonad and D) muscle+mantle. Measured concentrations are represented by coloured points according to the organ, and the linear model is given as the coloured line with the coloured area representing the 95% confidence interval. The equation, R² and p-value is given on each graph for initial and final times.

### 3.2 A seven-month in situ depuration (Dataset 2)

#### 3.2.1 Zone differences, depuration and body size relationship

The shell height (Supp. Figure S1A) did not significantly differ between the two zones (Wilcoxon test, p = 0.12, North: 8.7 *±* 1.6 cm, South: 8.4 *±* 1.8 cm). In contrast, flesh wet weight (Supp. Figure S1B) was significantly greater in the northern zone (47.7 *±* 20.9 *g*) than in the southern zone (41.3 *±* 19.0 *g*; p = 0.0013). The mean values and standard deviations of the condition index were similar between zones (North: 0.065 *±* 0.0083 *g cm*^3^, South: 0.061 *±* 0.0078 *g cm*^3^); while the difference was statistically significant (p = 10*^−^*^5^). This result reflects the large sample size and corresponds to a moderate size effect (Cohen’s d = 0.5), indicating that the mean condition index in the northern zone is approximately half a standard deviation higher than in the southern zone. The domoic acid concentrations were significantly greater in the northern zone (p = 0.0018), with a concentration of 18.1 *±* 11.4 *mg kg^−^*^1^ (range: 0.9-52.6 *mg kg^−^*^1^), compared to 14.2 *±* 10.6 *mg kg^−^*^1^ (range: 0.5-45.4 *mg kg^−^*^1^) in the southern zone. Consequently, the two zones were considered independently. Shell height and logarithmically-transformed domoic acid concentration were positively correlated (**Figure 6**), represented by 65% of the variance for the northern zone (**Figure 6A**) and 76% for the southern zone (**Figure 6B**).

**Figure 6:**
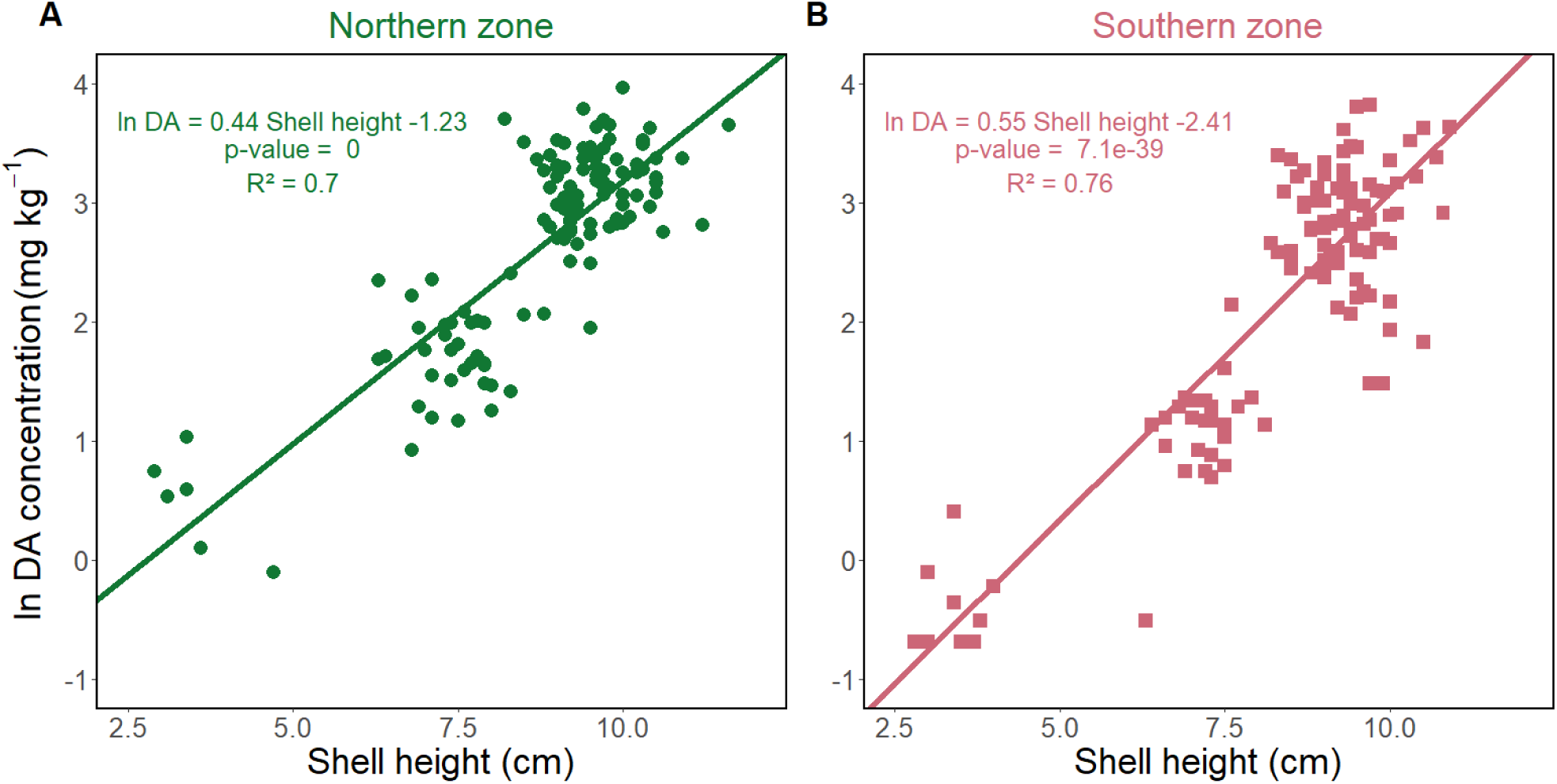
Domoic acid (DA) concentration in total flesh (*mg kg^−^*^1^) transformed by logarithm, according to shell height (*cm*) for A) the northern zone and B) the southern zone. The lines represent the linear model with equation, p-value and adjusted *R*^2^.

#### 3.2.2 Back-calculation of domoic acid concentrations at contamination

Based on the dataset 2 and on the REPHYTOX monitoring programme, the depuration rate was estimated at 0.008 *d^−^*^1^ for the northern zone (**Figure 7A**) and 0.009 *d^−^*^1^ for the southern zone (**Figure 7B**).

**Figure 7:**
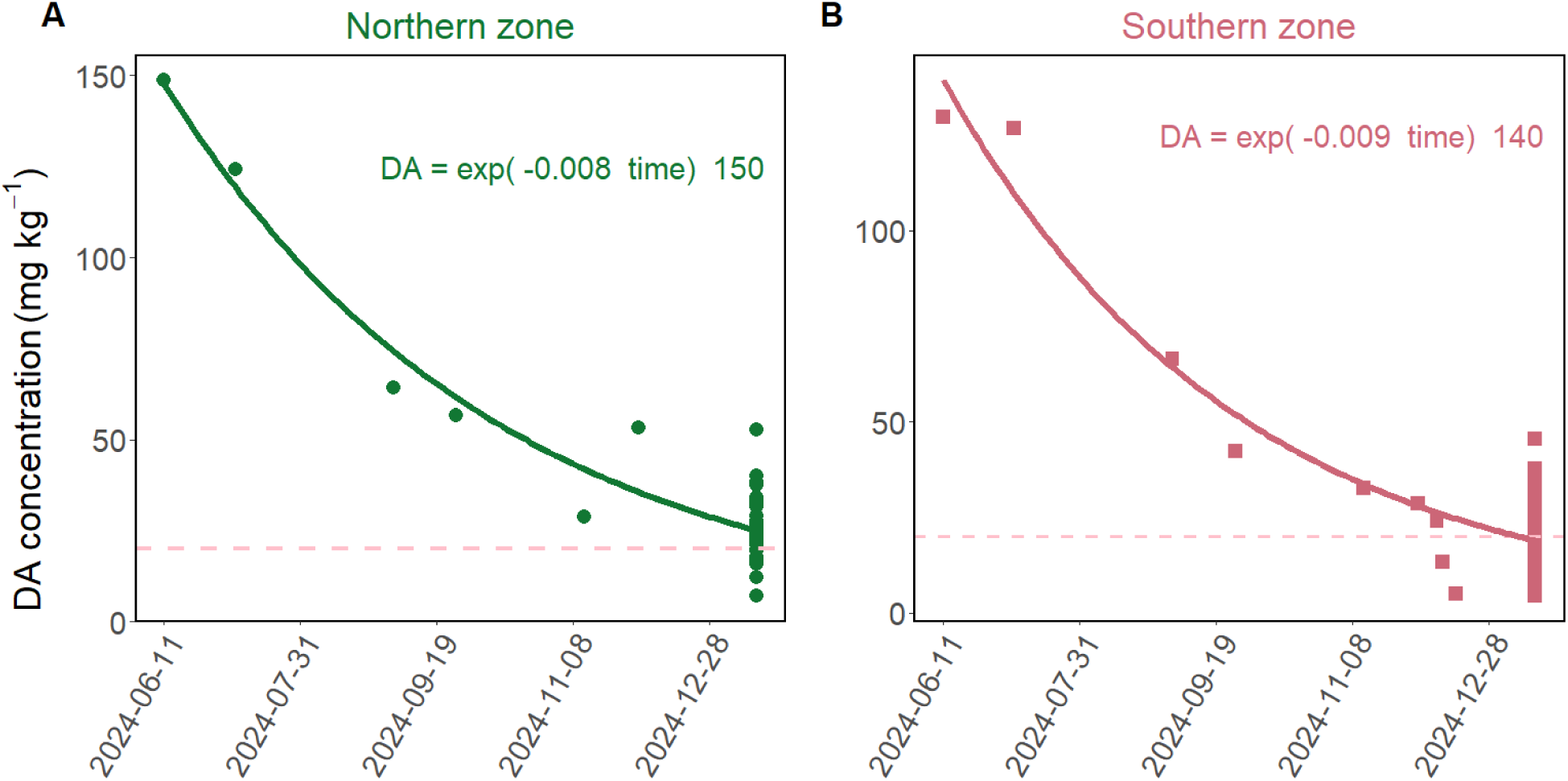
Domoic acid (DA) concentration (*mg kg^−^*^1^) in *P. maximus* exposed to a natural bloom of *P. australis* in April 2024 in the Bay of Brest according to date of sampling (“year-month-day”), for A) the northern zone and B) the southern zone. The individual dots from June to December 2024 correspond to measurement in the REPHYTOX monitoring (one pool of 10 individuals per sampled date) for the “039 - Rade de Brest - 039-S-281 Rade de Brest - Nord” REPHYTOX location, and the “039 - Rade de Brest - 039-S-282 Rade de Brest - Sud” REPHYTOX location. The dots in the last date are the individual measurements taken from sampling during this study. The plain line represents the estimation of the depuration dynamics, based on the exponential decay model using the “nls” function in R. The resulting equation is given on the graphs. The regulatory limit of 20 *mg kg^−^*^1^ is indicated by a pink dashed line.

The three scenarios to back-calculate the domoic acid concentration in June 2024 from January 2025 were applied to each individual. When applying dilution by growth (**Figure 8A, B**) only, domoic acid concentrations decrease according to shell height. However, this correlation was not significant for any of the two zones (p > 0.1). The average domoic acid concentrations for individuals above the minimal commercial length of 10.5 *cm* (shell height above 8.5 *cm*) were 36.7 *±* 13.0 *mg kg^−^*^1^ for the North and 24.4 *±* 12.2 *mg kg^−^*^1^ for the South (**Figure 8A, B**). A positive correlation (p < 0.001) between domoic acid concentrations and shell height was found with the scenario considering the estimated depuration rate only (**Figure 8C, D**). The correlation represented 45% and 40% of the variance for the north and south zones, respectively. The average back-calculated domoic acid concentrations for individuals above the minimal commercial length were 141.4 *±* 48.8 *mg kg^−^*^1^ for the North and 137.6 *±* 64.3 *mg kg^−^*^1^ for the South (**Figure 8C, D**). By applying both dilution by growth and estimated depuration rate (**Figure 8E, F**), the domoic acid concentrations and shell height were negatively but not significantly correlated. The average domoic acid concentrations for individuals above the minimal commercial length were 209.7 *±* 74.1 *mg kg^−^*^1^ for the North and 173.8 *±* 86.7 *mg kg^−^*^1^ for the South (**Figure 8E, F**). Therefore, the back-calculated average concentrations, resulting from the estimated depuration rate only, were the closest to the observed domoic acid concentrations in June 2024: 148.8 *mg kg^−^*^1^ in the North and 129.6 *mg kg^−^*^1^ in the South.

**Figure 8:**
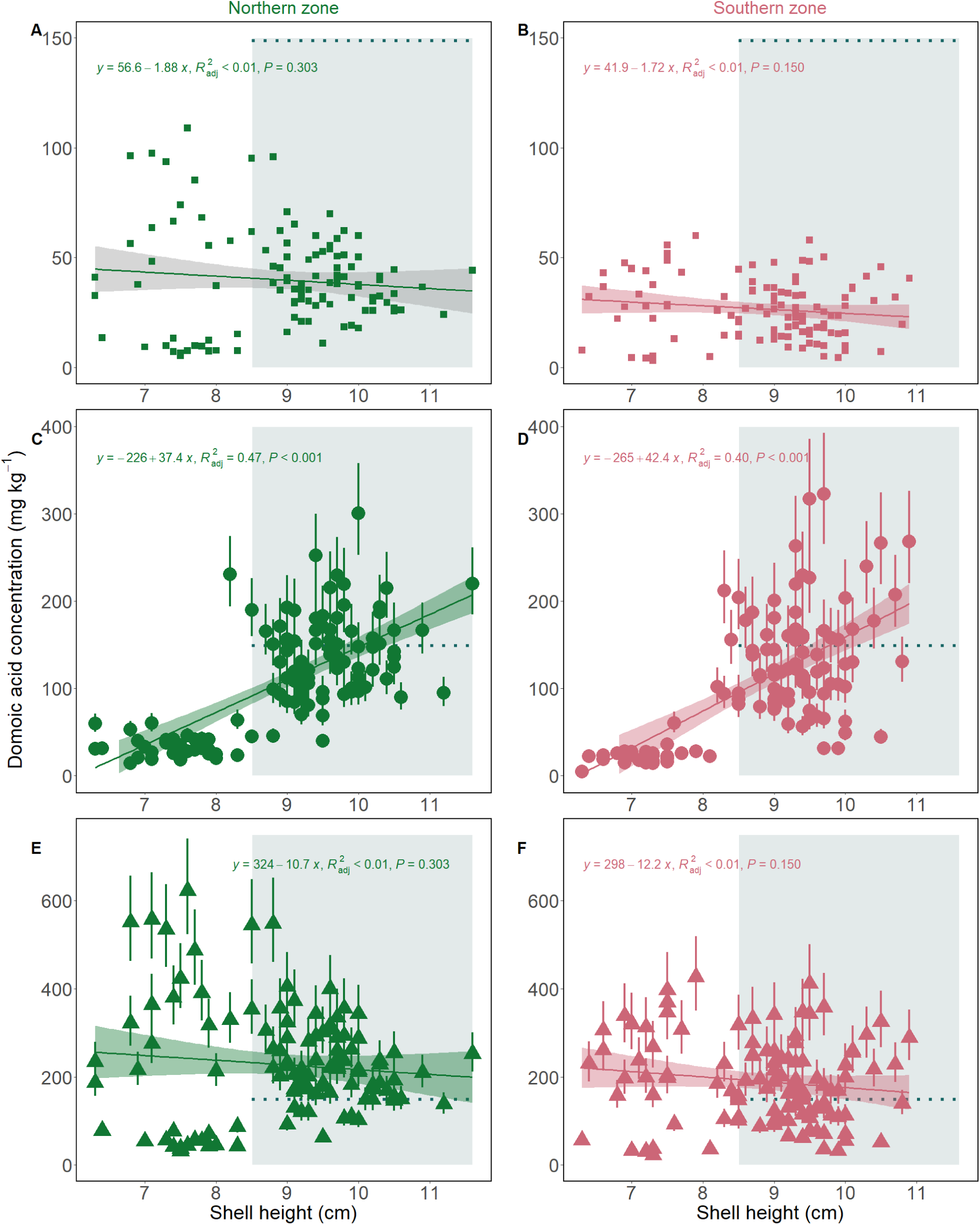
Simulated domoic acid concentration (*mg kg^−^*^1^) according to shell height for scenarios of dilution only (A-B), depuration only (C-D) and dilution + depuration (E-F), for northern zone (left panels; A, C, E, green points) and southern zone (right panels, B, D, F, pink points). For depuration and both processes scenarios, the points represent the simulations with median depuration rate and the bars show the simulations with slow and fast depuration rates. The dotted blue lines represent the measured domoic acid concentration in June 2024 by the REPHYTOX monitoring programme, thus only for individuals with commercial length (shell height above 8.5 *cm*) represented with the light blue square. The coloured line (green and pink) represent the linear model with its 95% confidence interval as the coloured areas. The equation, adjusted R² value, p-value and number of data used for the model are given on top of each graph.

## 4 Discussion

The objective of this study was to provide novel insights into the potential correlation between depuration of domoic acid by *P. maximus* and individual physiological traits, particularly body size. The first aim was to study the proportion of domoic acid in the different organs and how they vary throughout the depuration period. This study further focused on assessing the dependency of depuration on body size. Finally, the potential effects of growth on the depuration dynamics were investigated. Quantifying these correlations allows a better understanding of the mechanisms involved in toxin accumulation and depuration, thus providing useful information to model these processes. These combined results will provide valuable tools for scallop fishery management facing domoic acid contamination.

### 4.1 Proportion of domoic acid in organs and potential transfer

The digestive gland contained the highest concentrations and proportions of domoic acid, holding 97-98% of the total domoic acid burden both during the initial contamination phase and after a two-month depuration period (dataset 1). This finding aligns with previous studies (Blanco et al., 2002; Bogan et al., 2007). Domoic acid was detected in each organ, which is consistent with the observations of García-Corona et al. (2022) using immunohistochemistry. In this study, the muscle+mantle complex contained 2% of the total toxin burden, while the gonad accounted for 0.4 to 0.6%, which are comparable to the proportions reported for scallops (Campbell et al., 2001). However, Campbell et al. (2001) analysed the adductor muscle separately and grouped the digestive gland and mantle, resulting in muscle burden proportion of 0.17%. Although the gonad contributes to less than 1% to the overall toxin burden, its domoic acid concentration can still exceed the regulatory limit. Indeed, at the final time of the experimental depuration monitoring, the domoic acid concentrations in the gonads were above 20 *mg kg^−^*^1^ for a few individuals. This observation is consistent with previous findings (Blanco et al., 2002), which recorded proportions of around 2% for the gonads and kidneys. From an operational perspective, these results highlight the need to monitor domoic acid concentrations in separated organs, especially muscles and gonads, before the sale of shucked scallops as they may contain domoic acid concentrations above the regulatory limit.

Another major finding is that the distribution of domoic acid quantity and concentration can vary among organs and over time. Over a 2-month depuration period, the toxin concentrations decreased in both the digestive gland and the muscle+mantle complex, but increased in the gonads. Meanwhile, domoic acid burden decreased in the digestive gland, in the muscle+mantle, and slightly in the gonads. Álvarez et al. (2020) suggested that in *A. purpuratus*, domoic acid is transferred to the gonads. In this study, this hypothesis cannot be ruled out, but cannot be demonstrated either, as the constant domoic acid burden in the gonads, despite their weight loss, could either be associated to a transfer from other organs, or to differential depuration rates among organs, with gonads mostly retaining the toxin. Consequently, our findings do not clearly support a net transfer of domoic acid from other organs to the gonads.

The pattern of domoic acid distribution in organs may depend on several factors: the timing of the measurement relative to contamination and depuration phases, environmental conditions and the physiological conditions of the organism. First, regarding timing, Blanco et al. (2002) found no net toxin transfer and suggested that such transfer may only occur during the accumulation phase, which was not monitored. In the first experiment of this study (dataset 1), the initial sampling coincided with the contamination phase during a *P. australis* bloom, while the second sample was taken two months after. Consequently, this study only assessed the potential variation in the proportion of domoic acid burden in each organ between the contamination and the beginning of the depuration phase. Subsequent experiments should focus on the accumulation phase. The pattern of domoic acid distribution in organs may also depend on the environmental conditions, particularly food availability, which likely differs between in situ monitoring and laboratory experiments. During the experiment (dataset 1), the overall condition index and wet weights of the gonad and digestive gland decreased over time, suggesting a decline in scallop condition. Scallops typically use energy reserves stored in the digestive gland (Comely, 1974; Pazos et al., 1997). The concurrent weight loss in both the digestive gland and the gonads suggests that the individuals were partially starved and reliant on internal reserves, as observed in *Mimachlamys varia* (Régnier-Brisson et al., 2024). In scallops, energy mobilisation during starvation can be supported by gamete atresia (Le Pennec et al., 1991). It is therefore important to consider both the toxin quantity and the variations in organ weight, which reflect the overall conditions of the organism. Additionally, the natural physiological cycle of the individual, particularly the reproductive cycle, may affect observations of potential toxin transfer, as it affects gonad growth and reduction. In this study, gonad weight decreased after 60 days, whereas Blanco et al. (2002) observed an initial increase after 100 days, followed by a 50% decrease after 300 days. These findings highlight the importance of considering the gonadal cycle in both laboratory and natural settings.

To better assess possible toxin redistribution, based on the results of this study, one recommendation would be to collect individual samples at regular intervals during natural depuration and to quantify domoic acid concentrations in specific organs to track temporal changes. If the transfer of domoic acid to the gonads is confirmed, scallops could be authorised for market after the shucking process if they will be sold without their gonads.

### 4.2 Correlations between domoic acid concentration and body size

The correlation between domoic acid concentration and body size varied over the depuration period. After the contamination and following the two-month experimental depuration (dataset 1), the toxin levels were negatively correlated with the individual size: smaller scallops exhibited higher toxin concentrations. However, after the seven-month in situ depuration (dataset 2) which included a broader size range (2.5-12 *cm* shell height), larger scallops showed higher toxin concentrations, resulting in a positive correlation. This shift suggests that smaller scallops initially accumulate more domoic acid and depurate faster, while larger individuals retain the toxin over extended depuration periods.

A similar pattern was observed at the organ level. During the two-month experimental depuration (dataset 1), the domoic acid concentrations in the digestive gland and muscle+mantle were negatively correlated with shell height. In contrast, a positive correlation with size emerged in the gonads after the same period. These findings are partially consistent with those of Bogan et al. (2007) for *P. maximus* exposed to *Pseudo-nitzschia*, who reported a negative relationship between domoic acid and size at contamination and a positive one after 9 months of depuration. Their analysis, however, was limited to the digestive gland. Although domoic acid concentrations were not measured in the digestive gland after the seven-month in situ depuration (dataset 2), the fact that this organ accounts for 97% of the total toxin burden suggests that whole-body patterns reflect changes in the digestive gland.

Comparable size-dependent patterns have been documented in other bivalves. For example, oysters showed a negative relationship between their size and their domoic acid concentration during accumulation (Mafra et al., 2010), while no clear correlation has been identified in mussels (Novaczek et al., 1992). Additionally, smaller oysters depurate domoic acid more rapidly (Mafra et al., 2010). In scallops, higher toxin levels in smaller individuals have been attributed to higher uptake rates, and faster depuration in small scallops may explain the inversion of size-concentration correlation during prolonged depuration (Bogan et al., 2007). In this study, the estimated depuration rates over the seven-month period aligned with previously reported values for *P. maximus* (Blanco et al., 2002, 2006). The results from dataset 1 strongly indicate that, during a *P. australis* bloom, smaller scallops tend to accumulate higher toxin levels.

For sanitary monitoring programmes, these size-related differences have practical implications. In France, samples consist of a pool of 10 individuals above commercial size (shell length *≥* 10.5 *cm*), which smooths out the inter-individual variability and may not fully capture the range of toxin concentrations across all exploitable sizes. To address this, it would be interesting to either require samples from a defined broad size interval or systematically report the size distribution of each sample, along with a confidence interval. This approach would provide a more representative assessment of the inter-individual variability in domoic acid concentrations.

### 4.3 The influence of growth on domoic acid depuration

Given the slow depuration of domoic acid in *P. maximus*, the depuration periods are usually very long. Dilution by growth may occur during these periods, reducing the concentration of toxins as body volume increases. While this effect has been discussed for *P. maximus* (Blanco et al., 2002; Bogan et al., 2007), its quantitative contribution to depuration, to our knowledge, remained unexplored. Scallops exhibit von Bertalanffy-type growth (Mason, 1957; Buestel and Laurec, 1975): smaller individuals grow faster than larger ones. This suggests that smaller scallops are more effective at diluting their domoic acid burden during depuration. Similar patterns have been observed with other marine toxins, such as paralytic shellfish toxins in Atlantic surf clams, *Spisula solidissima* (Bricelj and Shumway, 1998) and ciguatoxins in fish, *Lagodon rhomboides* (Bennett and Robertson, 2021).

Simulating dilution by growth alone results in a slower decrease in toxin concentration than that observed. This process may help, however, to explain the observed inversion in the size-concentration correlation. Conversely, applying only the estimated depuration rate provides a good estimate of the average domoic acid concentration in *P. maximus* at the time of contamination. This is particularly true for scallops larger than the commercial size, for which data are available through the sanitary monitoring programme (REPHYTOX, 2023). Yet, this simulation does not account for the observed pattern in dataset 1, where smaller scallops exhibit higher domoic acid concentrations than larger ones. Unfortunately, the initial domoic acid concentrations for scallops in dataset 2 are unavailable, preventing validation of this back-calculation. For smaller scallops, integrating both the estimated depuration rate and dilution by growth better simulates the observed size-concentration inversion. Therefore, to accurately predict the decrease in toxin concentration, in terms of overall levels and body size patterns, both processes (dilution by growth and estimated depuration rate) must be considered.

The estimated depuration rate alone provides a robust average prediction of the decrease in toxin concentration, especially in commercial-size scallops, as already demonstrated in Le Moan et al. (2025). However, for smaller scallops, incorporating dilution by growth is essential. Two approaches could be considered for future modelling efforts to integrate dilution by growth alongside with estimated depuration rate. First, it is possible to adjust the depuration rate with a growth correction factor, as developed for ciguatoxin-contaminated fish (Bennett and Robertson, 2021). Second, a more integrative approach involves coupling an individual bioenergetic model with a toxicokinetic bioaccumulation model. Dynamic Energy Budget (DEB) models (Kooijman, 2010) have already been developed for *P. maximus* (Lavaud et al., 2014; Gourault et al., 2019) and for paralytic shellfish toxin accumulation in oysters (Pousse et al., 2019). Such a framework could provide insights into the mechanisms of domoic acid contamination in scallops.

## 5 Acknowledgements

The authors would like to thank the members of the Tinduff hatchery for the maintenance of the scallops during the first experiment, as well as the crew members of the Albert Lucas, vessel of IUEM and the scientific divers for the sampling of scallops in the Bay of Brest. We thank Juan Blanco for his useful and constructive comments on the manuscript, and Manon Bickert for proofreading it after the first round of peer review. Preprint version 3 of this article has been peer-reviewed and recommended by Peer Community In Ecotoxicology and Environmental Chemistry (https://doi.org/10.24072/pci.ecotoxenvchem.100455; Rodrigues, A. C. M. (2026) Size, organs and growth: key drivers of domoic acid depuration in king scallops. Peer Community in Ecotoxicology and Environmental Chemistry, 100455). We would like to thank our two reviewers for their valuable comments, which helped to improve the manuscript. We would also like to acknowledge the quality and rapidity of the reviews, as well as the recommendation, and PCI initiative.

## 9 Data and scripts

Data are available on: Le Moan Eline, Ambroziak Marie, Deléglise Margot, Vanmarldergem Jean, Derrien Amélie, Terre-Terrillon Aourégan, Breton Florian, Fabioux Caroline, Jean Fred, Flye-Sainte-Marie Jonathan, Hégaret Hélène (2026). Domoic acid concentration in king scallops associated with individual body size. SEANOE. https://doi.org/10.17882/111251 Scripts for analyses are available on: https://archive.softwareheritage.org/swh:1:dir:7a76109909405ffacf9f14dea985fb2424896a89

## Supplementary Materials for

**Table S1:** Precursor ion, product ions and MS/MS parameters used for the detection of domoic <u>acid.</u>

| Q1 Mass (Da) | Q3 Mass (Da) | Declustering Potential (V) | Dwell time (ms) | Collision Energy (eV) | Cell Exit Potential (V) |
| --- | --- | --- | --- | --- | --- |
| 312.1 | 161.1 | 82 | 200.00 | 33.00 | 12.00 |
| 312.1 | 193.1 | 82 | 200.00 | 25.00 | 16.00 |
| 312.1 | 220.1 | 82 | 200.00 | 28.00 | 18.00 |
| 312.1 | 248.1 | 82 | 200.00 | 24.00 | 12.00 |
| 312.1 | 266.1 | 82 | 200.00 | 22.00 | 14.00 |

**Figure S1:**
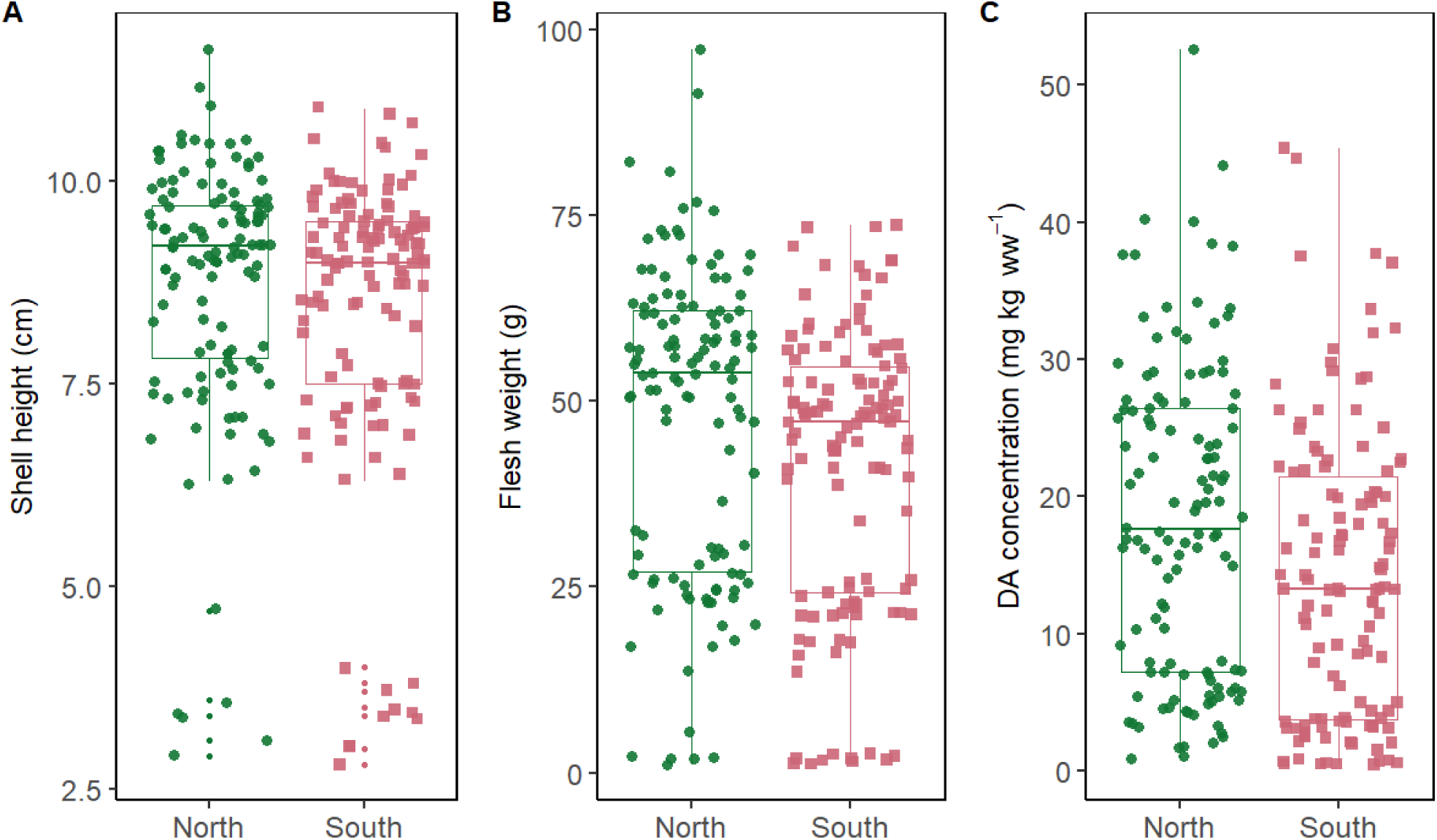
A) Shell height (*cm*), B) flesh wet weight (*g*) and C) domoic acid (DA) concentrations (*mg kg^−^*^1^) of individuals from each sampling zone.

